# The Role of the Temporal Pole in Temporal Lobe Epilepsy: A Diffusion Kurtosis Imaging Study

**DOI:** 10.1101/2021.10.07.463554

**Authors:** Loxlan W Kasa, Terry Peters, Seyed M. Mirsattari, Michael T. Jurkiewicz, Ali R. Khan, Roy A.M Haast

## Abstract

**Objective:** This study aims to evaluate the use of diffusion kurtosis imaging (DKI) to detect microstructural abnormalities within the temporal pole (TP) in temporal lobe epilepsy (TLE) patients.

**Methods:** DKI quantitative maps were obtained from fourteen lesional (MRI+) and ten non-lesional (MRI-) TLE patients, along with twenty-one healthy controls. This included mean (MK); radial (RK) and axial kurtosis (AK); mean diffusivity (MD) and axonal water fraction (AWF). Automated fiber quantification (AFQ) was used to quantify DKI measurements along the inferior longitudinal (ILF) and uncinate fasciculus (Unc). ILF and Unc tract profiles were compared between groups and tested for correlation with seizure duration. To characterize temporopolar cortex (TC) microstructure, DKI maps were sampled at varying depths from superficial white matter (WM) towards the pial surface. Each patient group was separated according to side ipsilateral to the epileptogenic temporal lobe and their AFQ results were used as input for statistical analyses.

**Results:** Significant differences were observed between MRI+ and controls (*p* < 0.005), towards the most anterior of ILF and Unc proximal to the TP of the left (not right) ipsilateral temporal lobe for MK, RK, AWK and MD. Noticeable differences were also observed mostly towards the TP for MK, RK and AWK in the MRI-group. DKI measurements correlated with seizure duration, mostly towards the anterior segments of the WM bundles. Stronger differences in MK, RK and AWF within the TC were observed in the MRI+ and noticeable differences (except for MD) in MRI-groups compared to controls.

**Significance:** The study demonstrates that DKI has potential to detect subtle microstructural alterations within the anterior segments of the ILF and Unc and the connected TC in TLE patients including MRI-subjects. This could aid our understanding of the extrahippocampal areas involved in seizure generation in TLE and might inform surgical planning, leading to better seizure outcomes.

## 1. INTRODUCTION

Temporal lobe epilepsy (TLE) is the most common form of medically intractable focal epilepsy in adults^1,2^. In most of these patients, the seizure onset zone lies within the mesial temporal lobe, which in 70% of the cases is induced by mesial temporal sclerosis (MTS), including hippocampal sclerosis^3^. Studies have shown, in clearly delineated MTS using both MRI (i.e., in MRI positive or ‘MRI+’ patients) and scalp EEG, that nearly 80% of patients are seizure-free after resective surgery^4^. Nevertheless, full delineation of pathological tissue can be challenging since seizures are not always exclusive to the hippocampus but may rather originate from extrahippocampal structures^3^. Thus, the epileptogenic zone may extend beyond the atrophic mesial temporal structures and explain long-term recurrence of seizures after selective resections^5^. A growing body of clinical investigation suggests that, among other extrahippocampal structures possibly involved in seizures, the temporal pole (TP, i.e., Brodmann’s Area [BA] 38 or anterior temporal lobe) could play an important and potentially underappreciated role in TLE^4,6^. The TP is connected to the three temporal gyri, while the two association fibers that terminate at the TP provide connections to the prefrontal cortex (i.e., uncinate fasciculus (Unc)) and amygdala and hippocampus (i.e., inferior longitudinal fasciculus (ILF)), consequently associating with several functions including memory^7,8^. Also, diffusion-weighted imaging (DWI) studies have consistently shown TLE to be a network disorder with underlying microstructural alterations in the temporal and extra-temporal white matter (WM) fiber bundles. Diffusion anomalies have also been detected in the cortical gray matter (GM) and superficial white matter (SWM, i.e., the WM area directly bordering the GM)^9–11^. Each of these separate studies found microstructural irregularities in the temporal pole, as depicted in the change of diffusion tensor imaging (DTI) parameters and decreased fiber density measured with neurite orientation dispersion and density imaging. Despite these promising findings using DWI, a substantial portion of TLE patients (∼30%) – often referred to as non-lesional (MRI-negative, or ‘MRI-’) patients – do not show lesions in their MRI scans, which can complicate the presurgical workup in these cases.

Although DTI is an elegant tool with relatively straightforward imaging requirements, a considerable number of studies (e.g., Tournier et al. 2011^12^ and references therein) have shown it to be inadequate for quantifying regions with complex fiber configurations (e.g., crossing fibers)^12^. As such, a technique called diffusion kurtosis imaging (DKI) was developed to address the DTI shortcomings that prevent it from accurately quantifying complex microstructure^13^. DKI enables the measurement of free diffusion (i.e., via its derived DTI metrics) and restricted diffusion within complex microstructure as it provides the common mean diffusivity (MD) and fractional anisotropy (FA) parameters, as well as different kurtosis measures, namely mean kurtosis (MK); radial kurtosis (RK); axial kurtosis (AK); and kurtosis fractional anisotropy (Kfa)^13,14^. These DKI-derived metrics have shown to be sensitive to WM network and GM abnormalities associated with TLE^15,16^. Furthermore, DKI can also be used to calculate specific microstructural compartments such as the ratio of axonal water content and the total water content per voxel, also known as axonal water fraction (AWF)^17,18^. Therefore, DKI could provide a complementary and more comprehensive characterization of diffusion in complex tissue environments, and potentially be more sensitive to diffusion anomalies in TLE patients compared to the more commonly employed DTI acquisition. However, the benefit of DKI for detecting subtle alterations in the microstructure of the TP in TLE, and MRI-in particular, has yet to be established.

The work described in this paper aims to evaluate the sensitivity of DKI to detect abnormalities at specific regions along the two association WM fiber bundles, ILF and Unc connected to the TP and the connected temporopolar cortex in MRI+ and MRI-TLE patients. We believe that a better understanding of the microstructural properties of the TP in TLE patients could improve the planning of resective surgeries and their outcomes.

## 2. MATERIALS AND METHODS

### 2.1. Subjects

Of the 24 TLE patients recruited in this study (9 females, mean age ± SD = 32 ± 10 years), 10 were considered MRI- (i.e., patients that do not show any signs of lesions in their structural scans, 4 female, 27 ± 6 years). The study was approved by the research and ethics board at Western University and informed consent was obtained from all patients and 23 healthy control subjects (14 female, 36 ± 15 years) prior to their recruitment in the study, following the Declaration of Helsinki. The patient cohort was selected based on the following inclusion criteria: (a) patients with a history of TLE; (b) had preoperative MRI, and (c) patients underwent comprehensive EEG studies to identify the site of their epileptogenic region. Thirteen patients had undergone temporal lobectomy (i.e., 6 right and 7 left hemisphere) and further investigation with post-surgical pathology confirmed the presence of MTS and gliosis in more than 75% of these patients. A detailed description of the demographic and clinical information for patients included in this study is provided in Table 1.

**Table 1.**
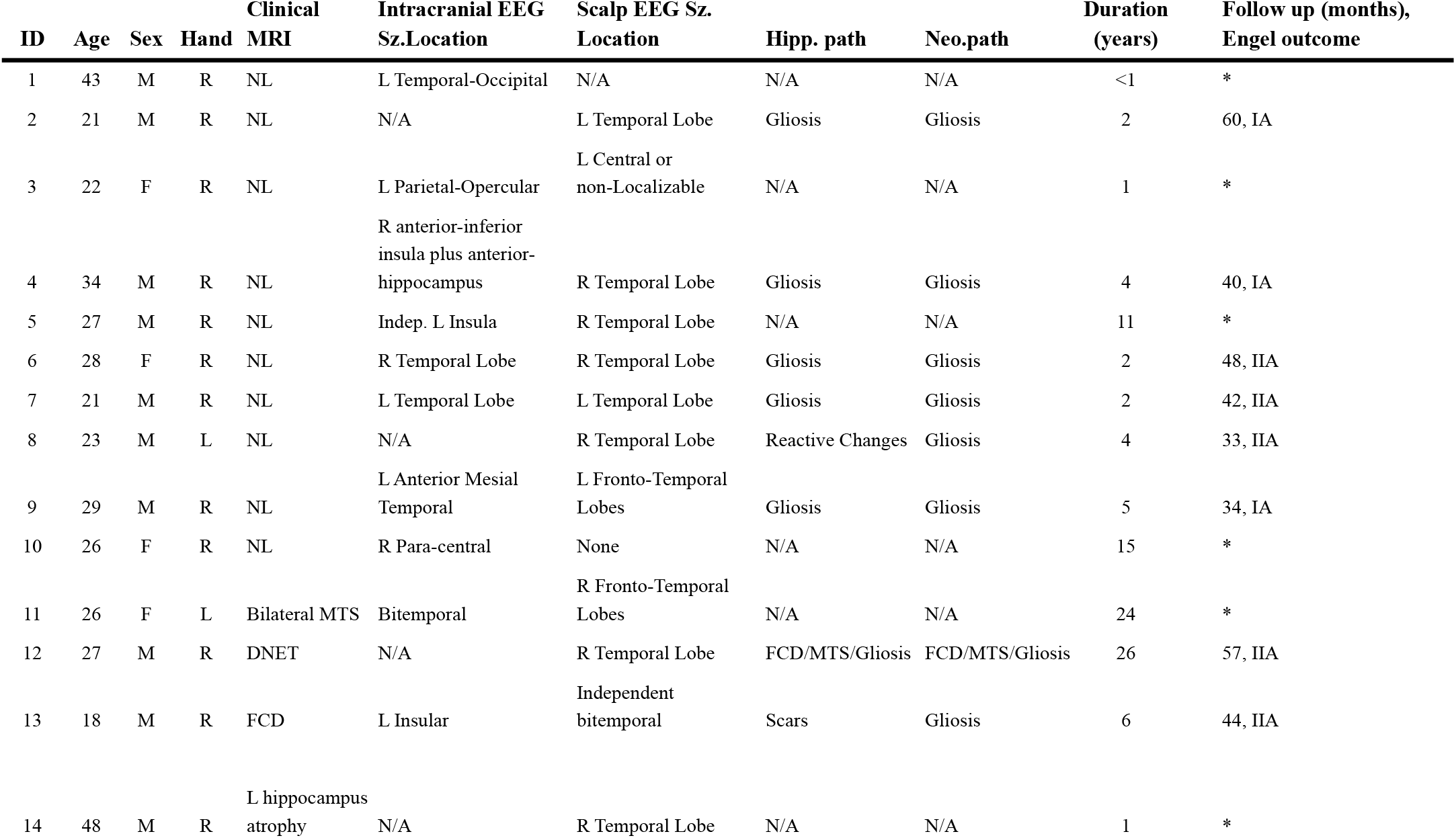
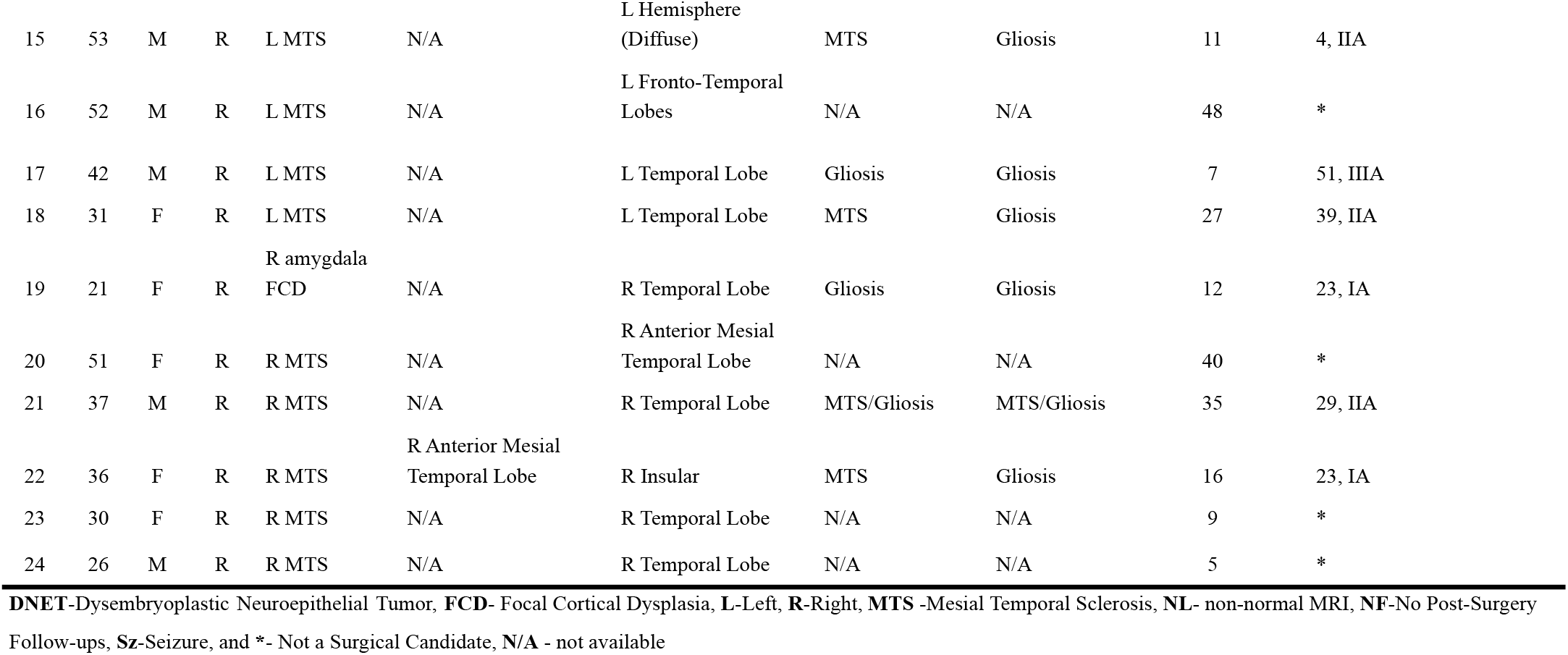
Clinical characteristics of patients with left and right temporal lobe epilepsy.

### 2.2. MRI acquisition and processing

All subjects were scanned using a 3T MRI system (Siemens Prisma, Erlangen Germany) with a 32-channel head coil. The scanning protocol included the acquisition of structural images using a magnetization-prepared rapid acquisition with gradient echo (MPRAGE) sequence (repetition time/echo time TR/TE = 5000/2.98ms, 700ms TI, FOV = 256×256mm^2^, 1mm isotropic voxel size). In addition, a multiband echo-planar imaging (EPI) sequence with acceleration factor = 3 was used to acquire diffusion-weighted images (DWI). The DWI acquisition includes b=0, 1300, 2600 s/mm^2^, 130 diffusion-encoding directions acquired twice with left-right, right-left phase encoding directions, TR/TE = 2800/66.80ms, FOV =224 × 224mm^2^, and 2mm isotropic voxel size. The acquired diffusion-weighted images were corrected for EPI readout and eddy current distortions using *topup*^*19*^ and *eddy*^*20*^ from FSL^21^. Correction of Gibbs’ ringing artifact was performed by determining optimal sub-voxel shifts within the neighborhood of sharp edges in the image^22^, while noise reduction was achieved by separating the signal from the noise in the image via local noise estimation (MRtrix3’s *dwidenoise*)^23,24^.

### 2.3. Calculations of DKI parameters

The preprocessed DWIs were used as input to the open-source Diffusion Kurtosis Estimator (DKE) software package to calculate the DKI parameters (MK, AK, RK, Kfa) including MD and FA derived from DKI^25^. In addition, we modelled the AWF metric from the kurtosis tensor using the DKE software^18^.

### 2.4 Automated white-matter fiber quantification

The automated fiber-tract quantification (AFQ) software^26^ was used to quantify the WM fiber bundles of interest (i.e., ILF and Unc). The basic AFQ four-step procedure was executed: (i) fiber tractography, (ii) fiber tract segmentation using two waypoint regions of interest (ROIs), (iii) fiber tract refinement via a probabilistic fiber tract atlas, and (iv) sampling diffusion measurements at 100 equidistant points along the tract length between two waypoints. To accommodate complex fiber configurations (i.e., fiber crossings) we modified the steps (i) and (ii) of the AFQ workflow, and each of these steps is described in the Supplementary Materials (sections S1.1 - S1.3).

### 2.5. Correlation between combined DKI metrics and seizure duration

Since DKI quantifies both the Gaussian component of diffusion (i.e., DTI metrics) and the non-Gaussian component (i.e., DKI metrics), we performed a combined DKI quantitative analysis (e.g., MD + MK). In addition, we incorporated the patient’s seizure duration (i.e., time between age of onset and age at scan) to investigate its correlation with different combinations of the DKI metrics.

### 2.6. Tract-based cortical analysis

To quantify diffusion profiles along the ILF and Unc to pial GM axis – referred to forthwith as tract-based cortical analysis (TCA) – we mapped all the individual subject’s DKI maps onto a series of cortical surfaces. First, a surface-based representation of the cortex was constructed using FreeSurfer’s *recon-all* pipeline and the MPRAGE T_1_w volume. The resulting WM surface was then used to obtain the additional cortical surfaces using FreeSurfer’s *mris_expand* function. These were positioned at different depth fractions based on the estimated local (i.e, vertex-wise) cortical thickness, starting with a ‘superficial’ WM surface (−50% of cortical thickness with respect to WM-GM boundary) up to the pial GM (+90% of cortical thickness) with steps of 10%, resulting in a total of 15 surfaces while maintaining smoothness and preventing self-intersection. FreeSurfer’s *mri_vol2surf* function was then used to project the DKI maps in anatomical space onto each of the generated surfaces. Due to limiting voxel size and partial volume effects, only values up to 50% (i.e., mid-GM) are considered for interpretation. Finally, vertex-wise DKI data were averaged within the rostral middle frontal (RMF) and temporal pole cortical regions (defined by FreeSurfer’s cortical parcellation) at each depth fraction per subject to allow comparison of profiles across groups. The RMF was included to serve as a baseline, as we expected little to no effects relating to TLE in this area. Similar to the WM analysis, for group-wise statistical analyses, each patient group (i.e., MRI+ and MRI-) was separated according to side ipsilateral to the epileptogenic temporal lobe.

### 2.7. Statistical analysis

The Permutation Analysis of Linear Models (PALM) toolbox was used to statistically assess group-wise differences in terms of WM profiles for each of the individual DKI-derived and WM distance maps^27^. A total of N=5000 permutations was used together with a cluster-wise *t*-statistic threshold of 3.1, while correcting for multiple comparisons (i.e., locations along the bundle) using the familywise error (FWE, q-FWE = 0.05), as well as age and sex effects. Output p-values were saved as -log10(*p*) for visualization. The *statsmodels* (v0.12.2) Python package was used for the comparison of WM profile shapes across groups using one-way analysis of variance (ANOVA), and between bundles (within-subjects) using repeated-measures ANOVA. As for the WM bundles, age and sex were accounted for by including them in the statistical models. Similarly, ANOVA testing was used for contrasting WM bundle summary scores between MRI+ and MRI-as well as diffusion parameter maps, while linear regression was used for testing the correlation with disease duration.

## 3. RESULTS

### 3.1 White matter quantitative profiling

DKI quantitative profiles and corresponding z-scores (shown as heat maps) for the MRI+ subjects, compared to the healthy controls, are shown in Figure 1A and 1C respectively. Ipsilateral to the seizure focus, significant differences were observed between MRI+ and controls (*p*<0.005) towards the most anterior end (i.e., position 100) of the left ILF for MK, RK, AK, AWF, and MD. A comparable, but slightly weaker, pattern was observed in Unc for the left MRI+ patients, with significant changes observed closer to the temporopolar cortex area in MD (*p*<0.005) and MK, RK, AK, and AWF (*p*<0.05). For illustrative purposes, the corresponding *p*-values for the microstructural alterations based on MK were mapped along the respective 3D renderings of the fiber bundles (Figure 1B). As such, it can be observed that MK for the left ILF (*p*<0.005) and Unc (*p*<0.05), differ most near the ipsilateral, anterior segments of the bundles (i.e., proximal to the temporopolar cortex). For the ipsilateral right temporal lobe, only AWF indicated possible microstructural alterations in the ILF (*p*<0.05), otherwise weak or insignificant differences were detected in the ipsilateral right temporal lobe of the MRI+ group.

**Figure 1.**
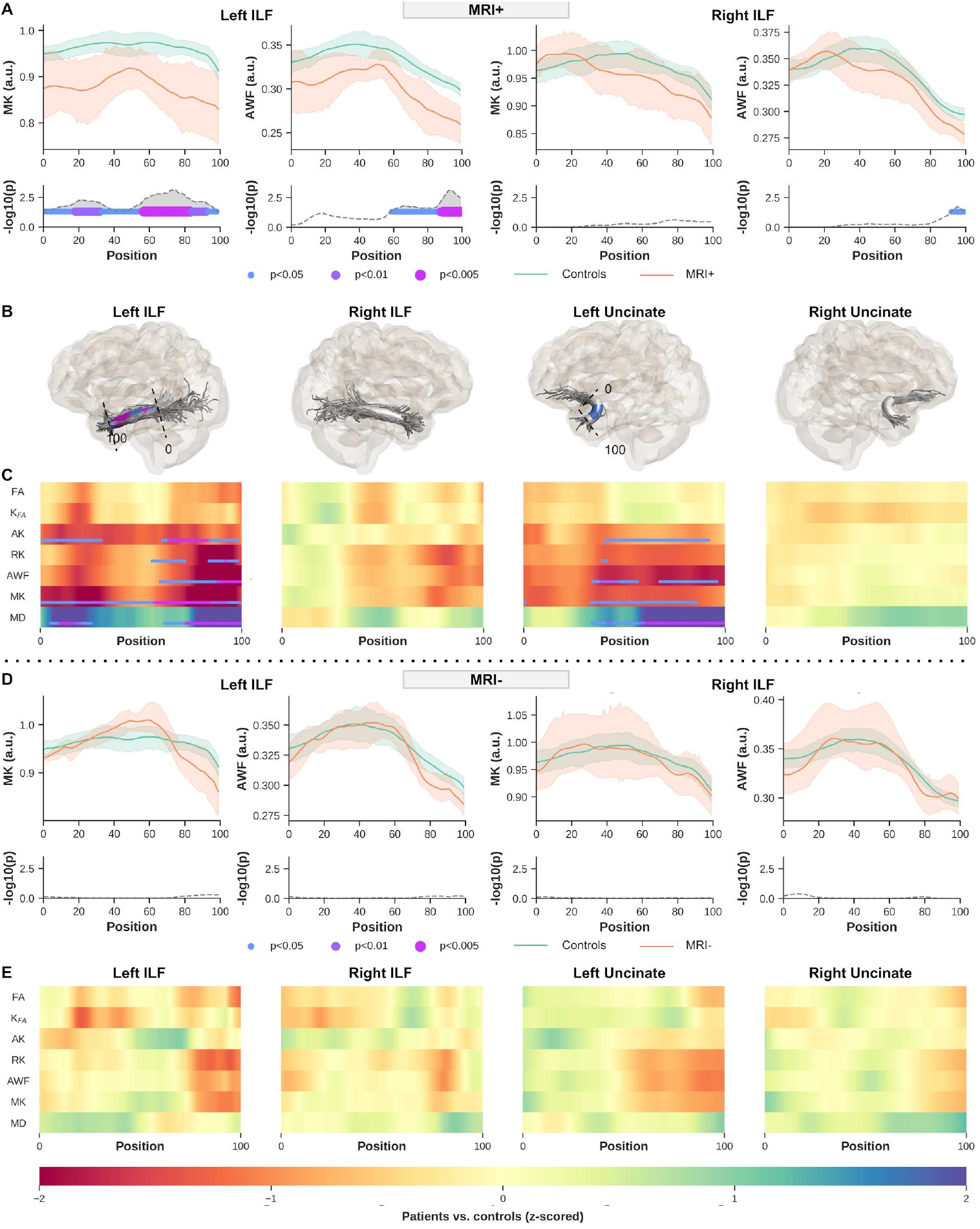
MRI+ patients vs controls (**A**-**C**). Showing only ILF MK and AWF profiles for ipsilateral left and right temporal lobe (**A**, top row) and corresponding *p*-values corrected for multiple comparisons using FWE, age, and sex (**A**, bottom row). To visualize the profile differences at corresponding anatomical locations along the bundles (0-100), the *p*-values are rendered onto the respective fiber bundles, showing here for MK only (**B**). Heat maps show z-scores for all DKI derived maps, for ILF and Unc (**C**). MRI-patients vs controls (**D**-**E**). Showing only ILF MK and AWF profiles for ipsilateral left and right temporal lobe (**D**, top row) and corrected for multiple comparisons using FWER, age, and sex *p*-values (**D**, bottom row). The heat maps show z-scores for all DKI maps including DTI MD and FA estimated with DKI, for ILF and Unc (**E**). Note that only results from the side ipsilateral to the epileptogenic temporal lobe are shown for each bundle.

For MRI-patients, noticeable differences were observed based on the DKI quantitative measurements, in particular MK, RK, and AWF, ipsilateral to the seizure focus (as confirmed with EEG), in line with the MRI+ patients and as indicated in Figure 1 (D-E). Although no significant differences were detected after multiple comparison corrections, indicators of possible changes along the WM bundles were mostly found at their most anterior parts in left TLE subjects, similar to what was observed with the MRI+ group (Figure 1E).

The profiles calculated from the distance maps from the two (i.e., MRI+ and MRI-) patient groups and the controls as shown in Figure 2, clearly indicate all DKI quantitative sampling were exclusively within the WM (i.e., negative distance, Figure 2A, top row). One thing to note, however, is the difference in the mean distance profiles from the WM to the WM-GM interface observed between the patient groups and the controls shown in Figure 2A, as well as demonstrated on the heat maps (Figure 2B), in particular ipsilateral left temporal lobe ILF and Unc (MRI+) and ipsilateral left temporal lobe ILF and right Unc for the MRI-group.

**Figure 2.**
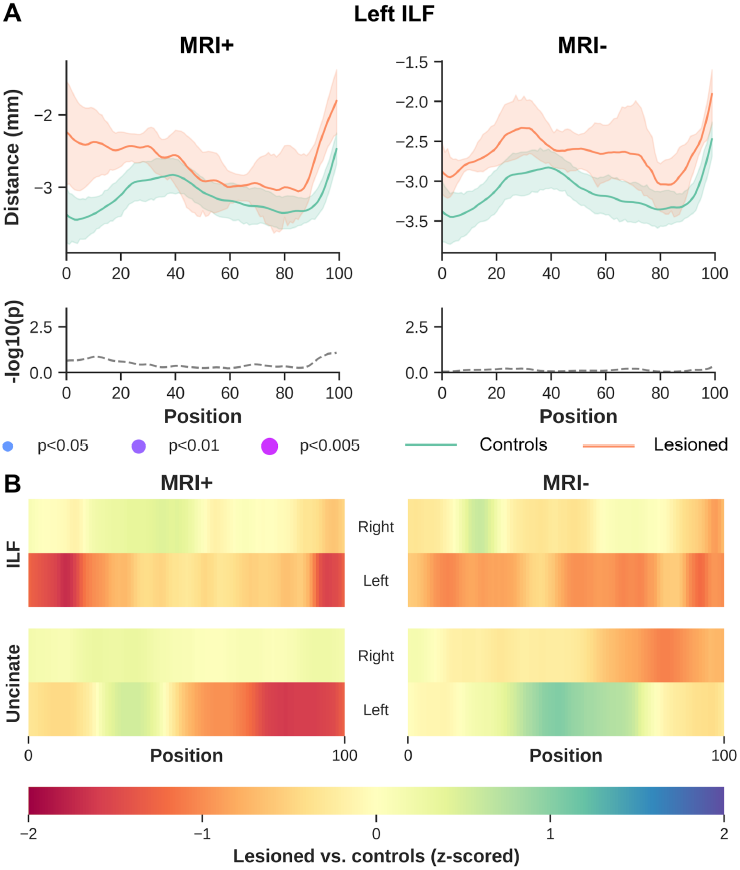
WM to GM distance profiles for MRI+ and MRI-ipsilateral left temporal lobe ILF (**A**, top row) and corrected for multiple comparisons using FWER, age, and sex *p*-values (**A**, bottom row). The heat maps show z-scores from the distance profiles for ipsilateral left and right temporal lobe ILF (**B**, top row) and for ipsilateral left and right temporal lobe Unc (**B**, bottom row).

### 3.2 White matter profile’s shape analysis

Geometrical properties of the quantitative profiles showed possible differences between the two bundles (ILF and Unc), as shown in Figure 3. To demonstrate the distribution of the average of the diffusion kurtosis along all diffusion directions^28^ we are showing MK only and AWF as a measure of possible changes in axonal density.

**Figure 3.**
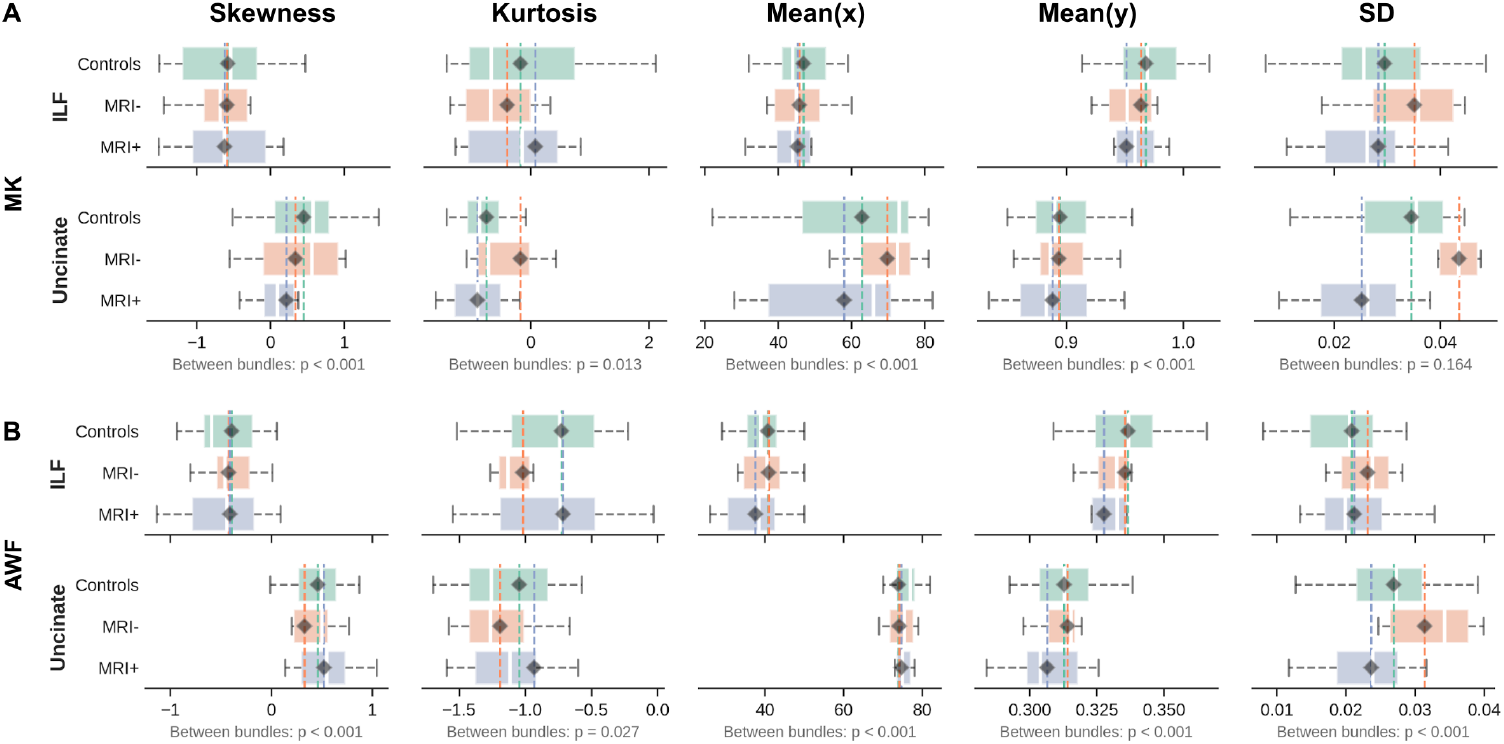
Shape analysis of DKI quantitative profiles from MK (**A**) and AWF (**B**) for the respective groups (y-axis, i.e., MRI+ in gray, MRI-in orange, and controls in green). The x-axes are separated with respect to the first four statistical moments; skewness, kurtosis, mean(x) — mean in the x-direction, mean(y) — mean in the y direction, and the standard deviation (SD) in the rightmost bin. Only ipsilateral left temporal lobe results are shown, as it appeared most affected based on the tract profile.

A more subtle and variable difference is observed between the patient groups compared to control’s profile shapes as depicted in their skew, kurt, mean(y), and mean(x) distributions. However, in both MRI+ and MRI-groups, the SD for the two WM bundles MK (Figure. 3A) and AWF (Figure. 3B) values show consistent dissimilarity compared to controls.

### 3.3. White matter correlational analyses

When combining the DTI metrics estimated with DKI (i.e., MK+MD and Kfa+FA), we observed significant (*p*<0.05) differences between the two patients groups (i.e., MRI+ vs MRI-) at the left anterior parts proximal to the temporopolar cortex (i.e., points 80-100) of the WM bundles within the ipsilateral temporal lobe (Figure 4A, top row). Higher, but not significant z-scores were also observed when combining MK+MD in ILF and Unc from the ipsilateral left temporal lobe, compared to the combination of Kfa+FA. We also observed a linear relationship with MK+MD in the anterior ILF with epilepsy duration as shown in (Figure 4A, bottom row), though based on visual judgment, a smaller change is noted with Kfa+FA values. A similar pattern is exhibited within the anterior Unc, showing a slight difference between the patient groups with longer seizure activity.

**Figure 4.**
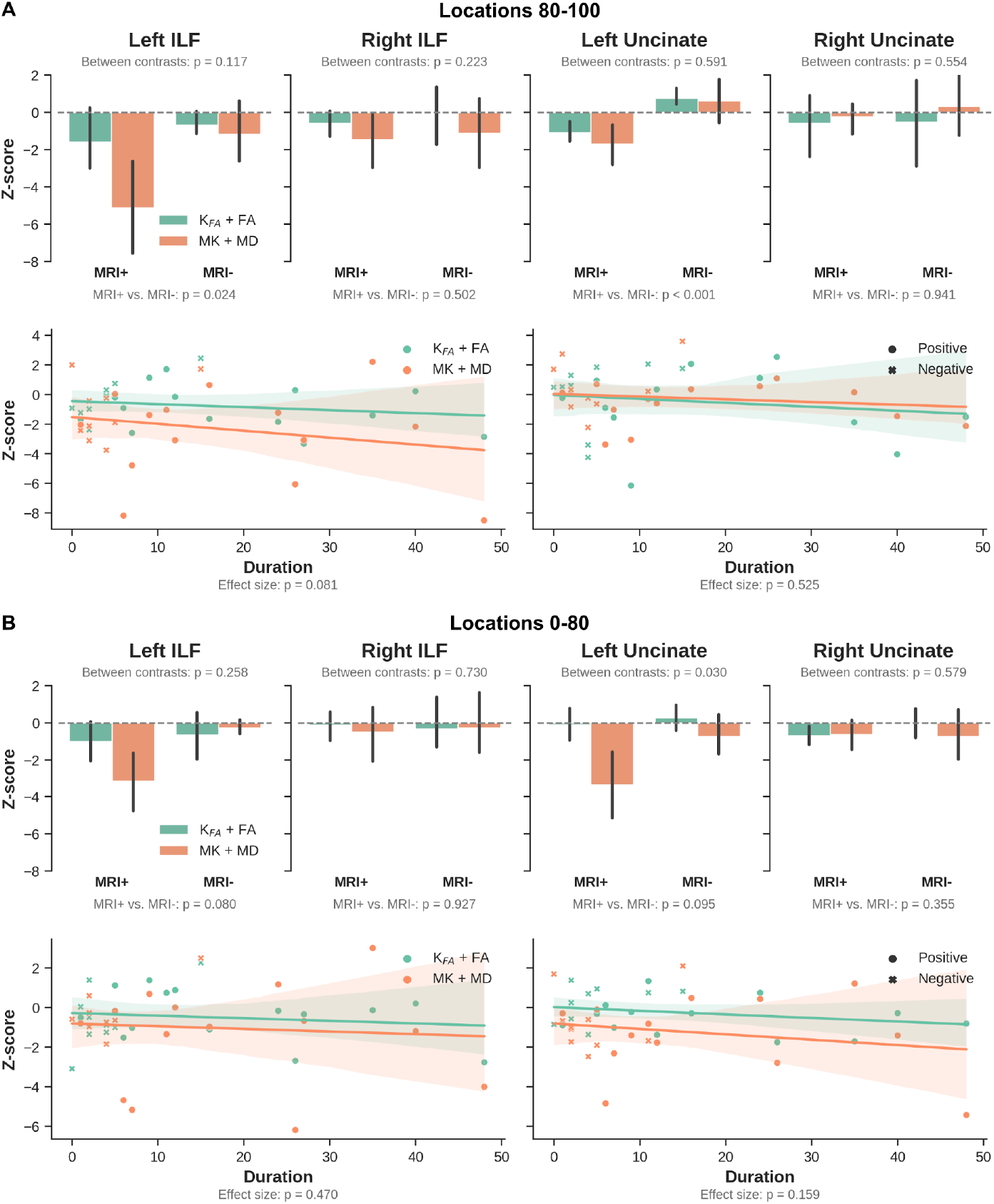
Combined DKI parameter analysis. The bar plots (top row, **A**) show z-scores (Kfa + FA, green) and (MK + MD, orange) for MRI+ and MRI-patients vs controls calculated within the ipsilateral temporal lobe for ILF (left columns) and Unc using values extracted from sampling points (80-100). The correlation of combined DKI quantitative values within ipsilateral left and right temporal lobe and seizure duration is given on the scatter plots (bottom row, **A**), with MRI+ (•) and MRI- (×). **B**, combined DKI parameter analysis, values extracted from sampling points (0-80). Between contrast (i.e., MK+MD vs Kfa+FA) and effect size shows FWER, age, and sex *p*-values.

No significant changes were detected between patient groups and WM bundles when averaging across sampling points 0-80. Nevertheless, in line with the observations for the anterior portions of the bundles, the strongest effects were observed for the ipsilateral left temporal lobe data. Here, MRI+ patients were characterized by the largest differences, driven primarily by the MK+MD parameters (Figure 4B, top row). A more comparable difference was noticed in the ipsilateral right temporal lobe between the patient groups’ MK+MD measurements. The DKI combined quantitative values show increased differences with persisting seizure in patients, with more distinct MK+MD changes in both ipsilateral temporal lobes (Figure 4B, bottom row).

### 3.4 Tract-based cortical analysis

We observed consistently strong differences at the temporopolar cortex tissue transition area (i.e., superficial WM towards WM-GM) between the controls and MRI+ group (Figure. 5B, top row) for the MK, RK, and AWF maps. In the MRI-group, we also see notable DKI differences, except for MD. In general, z-scores gradually change to zero while moving towards the pial surface. For both patient groups, almost no differences were observed at the RMF region (Figure. 5B, bottom row).

**Figure 5.**
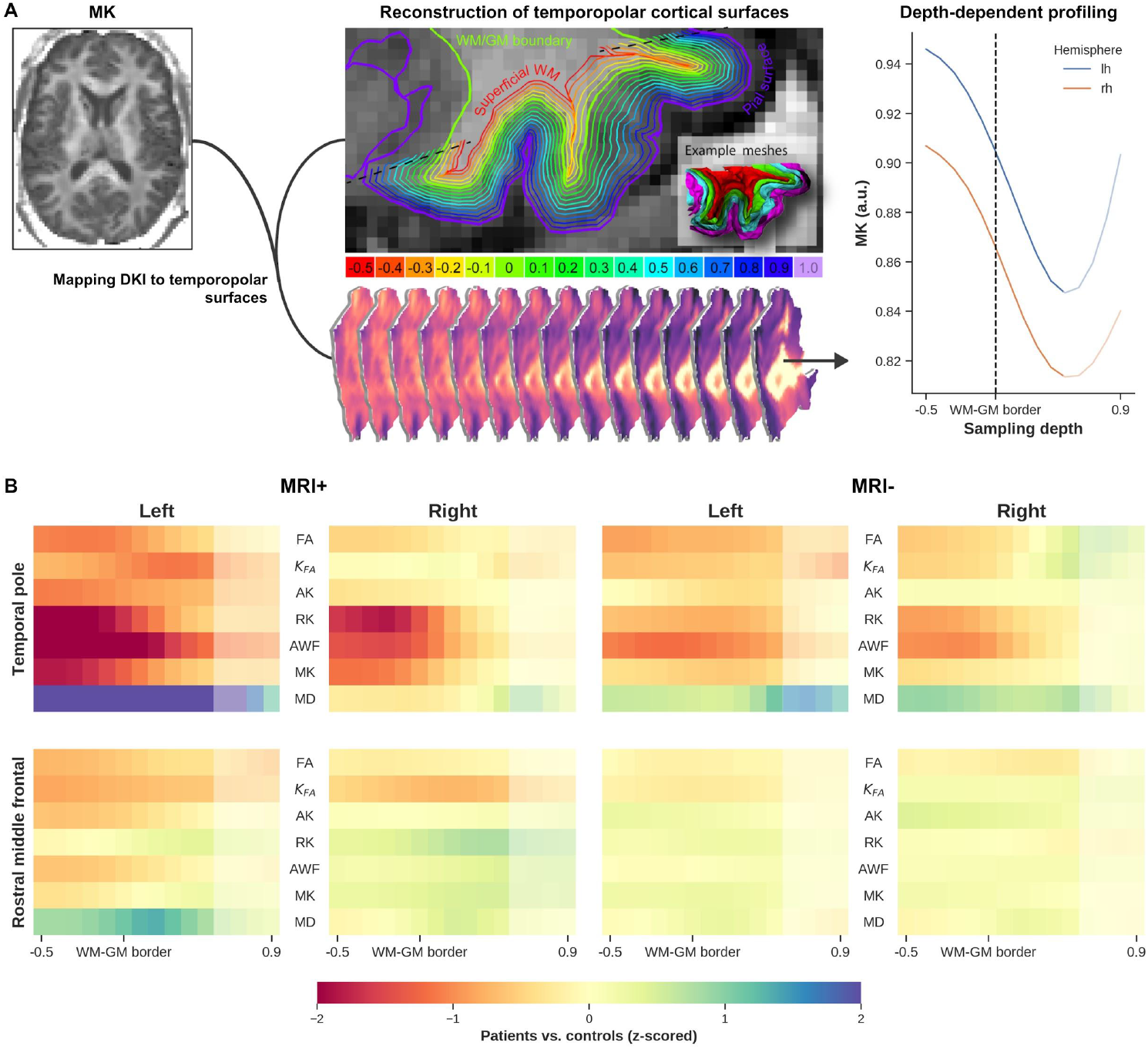
General workflow — cortical diffusion profiling (**A)**. The DKI quantitative values (e.g. MK, left **A**) are projected onto the temporopolar surfaces at varying depths, starting from the superficial WM (middle **A**, red) towards the pial surface (purple). Finally, quantitative measurements of the temporopolar cortex microstructure are extracted as depth-dependent profiles (right **A**). Comparison of cortical microstructure between patients and controls is shown in (**B**). Each heatmap shows z-scores for all DKI parameters extracted from the superficial WM (i.e., -0.5) towards the pial surface (0.9). The top row shows MRI+ patients vs controls (left columns) and MRI-patients vs controls (right columns) calculated within the temporopolar cortex of the side ipsilateral to the epileptogenic temporal lobe, similarly the bottom row presents rostral middle temporal. Note: The increased MK values towards the pial (**A**, line plot) show a possibility of cerebrospinal fluid (CSF) influence, so care should be taken when interpreting these results.

## 4. DISCUSSION

In this study, we combined the anatomically constrained tractography (ACT) using multishell constrained spherical deconvolution (CSD) (see supplementary data S1.1) with DKI to compare diffusional properties along (i) the ILF and Uncinate, two major WM fiber bundles connecting the temporal pole with other cortical regions, as well as (ii) the WM to the transition area of the temporal pole, between healthy controls and TLE patients. Most importantly, we found prominent diffusion profile differences closer to the anterior portions of both bundles within the side ipsilateral to the epileptogenic temporal lobe. In addition, diffusion anomalies were detected within the temporal pole cortex ipsilateral to the epileptogenic temporal lobe and were more pronounced in the DKI measurements of the TP compared to the reference RMF. This is the first study to combine ACT using multishell CSD to overcome limitations imposed by complex fiber configurations (e.g., crossing fibers), AFQ for tract profiling, and DKI to characterize tissue microstructure by taking into account non-Gaussian diffusion behavior. Furthermore, depth-dependent DKI measurements were extracted to highlight possible diffusion abnormalities from superficial WM towards the pial surface of the TP.

### 4.1 White matter quantitative profiling

Based on the diffusion profiles dissimilarities towards the anterior segments of the bundles, more prominent differences are noticeable for the ipsilateral left epileptogenic temporal lobe across the MRI+ patients, in particular with regards to MK, RK, AK (i.e., decrease), and MD (increase). These findings are in line with a previous study that looked at DKI- and DTI-based metrics along specific WM tracts of left TLE subjects^16^. Moreover, as a surrogate marker for axonal density, the decrease in AWF showed potential axonal degradation in the left side ipsilateral to the epileptogenic temporal lobe in MRI+ patients, which has been a common find in TLE patients with MTS^29–31^. We also observed a decrease in DKI parameters and an increase in DKI-based MD towards left TP in both of the bundles ipsilateral to seizure focus in the MRI-group. The differences in diffusion abnormalities between the left and right TLE patients, as categorized by seizure onset zone, agrees with a previous voxel-based study demonstrating widespread and prominent DTI abnormalities in patients with MTS and left hemispheric onset, while right MRI-patients exhibited no detectable changes^32^. This suggests that DKI could serve as a complementary approach to detect subtle changes in the WM fiber bundles connected to the TP. In addition, the observed reduction in the distance of the patients’ WM fiber bundles to the WM-GM boundary could be attributed to the presence of ectopic WM neurons (i.e., WM neurons in abnormal locations, mostly found in the subcortical region) which increase in density in brain specimens of TLE patients compared to controls^33^. Identifying abnormalities at specific locations along the ILF and Unc before surgery in TLE, particularly in MRI-subjects, could be used to guide the procedure by more completely identifying the seizure onset zone and the region to be resected, potentially resulting in improved outcomes. However, although the current findings demonstrate DKI’s potential to measure diffusion in patients, it strongly motivates further evaluation with a larger patient cohort.

### 4.2. White matter profiles shape analysis

Tract profile analysis of the MK and AWF values along WM bundles (i.e., between sampling points 0-100), indicated that ILF and Unc ipsilateral to seizure focus have a wider SD in the MRI-group (more pronounced in Unc). Here, SD represents the variation along the y-axis of the profiles, implying that MK and AWF appear to vary more along points 0-100 for the MRI-, but are more ‘flat’ for the MRI+ patients, compared to controls. This observation could correspond to the considerable evidence of dispersed diffusion abnormalities observed in MRI-patients compared to those with TLE patients having radiological evidence of MTS (i.e., MRI+)^29,34^. In addition, a smaller SD of the AWF data for the MRI+ patients could also suggest diffusion anomalies related to degradation of spatial specificity of microstructural properties along the WM pathways compared to the more variable changes exhibited with MRI-patients. Furthermore, between the bundles, Unc shows greater deviations than ILF in the MRI-patients, in concordance to a previous tract base study, which found a significant reduction in the Unc, that was more pronounced in the anterior temporal lobe^16^. These differences observed in the DKI parameter distribution, specifically SD, agree with the WM profiling analysis indicating potential diffusional changes due to patterns of neuronal loss and gliosis, particularly observed in patients with MTS^30^.

### 4.3 White matter correlational analysis

The complementary properties of the MK and MD (i.e., restricted vs. free diffusion) parameters were utilized by combining their respective z-scores. In agreement with the WM findings, a significant difference at the anterior portion of the bundles between the two patient types (MRI+ vs MRI-) was observed. MK+MD demonstrated potential microstructural changes in the ipsilateral left epileptogenic temporal lobe, where MRI+ showed stronger differences compared to MRI-group, especially towards the anterior segments of the two fiber bundles (i.e., from sampling point 80 right to the most anterior point 100). Similar, but milder diffusional changes were reflected in the MK+MD values within the more posterior segments (i.e., points 0-80) of the ILF and Unc, with noticeable differences with left TLE patients compared to controls. This goes along with a previous study that demonstrated a centrifugal decrease of DTI-based abnormalities as WM networks extend away from the epileptogenic temporal lobe^29,35^. Furthermore, there was a relationship between the MK+MD measurements and seizure duration in the two patient groups (MRI+ and MRI-). The detected progressive diffusion anomalies suggest gradual microstructural changes due to persistent seizure activities^36^. This pattern was also noted in a DTI-base study, with MD strongly correlating with seizure duration^37^. Although MK+MD showed clear differences compared to Kfa+FA, there were no significant differences between these combined DKI parameters (i.e., MK+MD vs Kfa+FA). Nevertheless, since MK and MD are average measurements along all diffusion weighting directions in restricted and free diffusion environments respectively, we expected MK+MD to provide a comprehensive characterization of diffusion properties depicting tissue integrity. On the other hand, FA depicts the preferred direction of diffusion and can reduce drastically in areas of crossing fibers or often in regions of coherent WM^38^. Therefore by combining FA and Kfa, complementary information regarding tissue microstructure could be derived^38^.

### 4.4 Tract-based cortical analysis

The present findings indicated that MK and RK can detect microstructure anomalies within the temporopolar cortex in MRI+, and to some extent within the MRI-patients compared to a reference region (i.e., RMF). Although TLE appears to be characterized by a network of abnormalities^15^, we expected RMF to have a minimal association with the temporopolar cortex changes due to seizure activities. Besides the observations related to MK and RK, the changes in AWF indicate potential neuronal loss. This could be due to seizure activities originating either at the temporopolar cortex or the medial temporal lobe structures (e.g. hippocampus)^3^. These findings support previous work which detected neurons and dendritic changes in the temporopolar cortex in TLE patients^10^. Parallel to this study and in support of our findings, several earlier imaging- and histology-based studies have also identified a reduction in neuronal density and gliosis within the superficial WM and temporopolar cortex^39,40^. Furthermore, the WM-GM boundary diffusion anomalies could be related to the blurring of this area, commonly attributed to various causes (e.g. developmental cortical abnormalities, gliosis, or myelin alterations)^33,41^. Based on the current results, DKI could serve as a complementary approach to detect subtle microstructural alterations within the temporopolar cortex. Understanding the role of TP in TLE is important to inform resection, which could help to minimize seizure recurrence due to insufficient resection^42^.

### 4.5 Limitations

The diffusion anomalous along the WM fiber bundles connecting TP and the connected temporopolar cortex detected with DKI measurements indicated DKI’s ability to quantify subtle alterations of the microstructure in TLE patients. However, future work may include larger cohort patients groups (MRI+ and MRI-) to validate these findings. The spatial resolution (2 mm isotropic) of the DWI used in this study is approximately equivalent to the cortical thickness^43^. Therefore, the current findings could still be affected by partial volume effects near CSF. As such, data closer to the pial surface (i.e., sampling depth > 0.5) were not taken into consideration. Nevertheless, as we were particularly interested in the WM-GM transition area, the current TCA is still valid to reveal trends in terms of microstructural differences in the TLE patients. Moreover, statistical results were corrected for age and sex to minimize their influence on the patient vs. control differences. However, age- and sex-matched samples could avoid any residual differences in diffusion characteristics due to aging or gender^44,45^.

## 5. CONCLUSIONS

The current study demonstrated that the combination of ACT multi-shell tracking, AFQ, and DKI could serve as a complementary approach to detect subtle microstructural alterations within the anterior segments of the two association WM fiber bundles connected to the temporal pole. In addition, depth-dependent DKI measurements could aid in uncovering diffusion abnormalities in the temporopolar cortex. Furthermore, since the DKI acquisition and precision have shown to be clinically feasible^46^, the methods developed in this study could be easily implemented in a clinical workflow. Finally, while the study was based on a limited patient cohort it provides solid preliminary data upon which to base a more comprehensive investigation. Identification of anomalies along the WM bundles segments and specific depths within the temporopolar cortex before surgery could help inform the planning of selective resection to improve outcome, particularly in MRI-patients.

## 5. ACKNOWLEDGEMENT

We would like to thank the patients and control participants who agreed to take part in this study. This work was supported by Canadian Institutes of Health Research (CIHR) Foundation, Natural Sciences and Engineering Research Council (NSERC) Discovery, the Canada First Research Excellence Fund, Brain Canada, and the Ontario Brain Institute Epilepsy Program (EpLink). Author R.A.M.H was supported by a BrainsCAN postdoctoral fellowship for this work.

