## Supplementary Methods for "The Role of the Temporal Pole in Temporal Lobe Epilepsy: A Diffusion Kurtosis Imaging Study"

Loxlan W Kasa, Terry Peters, Seyed M. Mirsattari, Michael T. Jurkiewicz, Ali R. Khan, Roy A.M Haast

### Supplementary Methods

#### S1.1. Fiber tracking and segmentation

We used the anatomically constrained tracking (ACT) algorithm[1] implemented in MRtrix3 software[2]. In ACT, the first step is to calculate a response function, which was estimated from the pre-processed DWIs using *dwi2response* (with the "dhollander" algorithm for multi-shell data)[3]. In the second step, the estimated response functions for individual DWIs were used to calculate the fiber orientation function (FOD) using *dwi2fod*. The FODs were separately calculated for the three tissue types (i.e., WM, GM, and cerebrospinal fluid (CSF)) using the multi-shell multi-tissue constrained spherical deconvolution (CSD) method (*msmt_csd*)[4]. To allow for group comparison, the subjects' WM FODs were input to the *population_template* function to generate an unbiased group average FOD template, to which the respective subject's FODs were warped. To deploy the ACT algorithm, additional anatomical information is required to guide the termination and acceptance/rejection criteria during fiber tracking in MRtrix3[1]. Individual subject's T1 images were segmented into five tissue types ('5TT'), namely: WM, subcortical GM, GM, and CSF, and the optional tissue type (i.e., pathological tissue, which was excluded here), using the *5ttgen fsl* command. The *5ttgen fsl* command uses the FSL FIRST and FAST segmentation functions to separate the four mandatory tissue types. To minimize tracking into the deep GM, we generated a WM-GM interface mask by inputting the 5TT segmented anatomical image into the *5ttgmwmi* command. The resulting WM-GM interface mask was used as the seed point for tracking. To further guide fiber tracking, extra parameters were supplied to the MRtrix *tckgen* function responsible for tractography, following the six criteria described by Smith et al. (2012)[1]. These parameters influence the streamlines in two ways: when to terminate them and when they are either accepted or rejected based on their biological plausibility. Since tracking was seeded in the WM-GM interface, two waypoints manually created for the bundles ILF and Unc were input as part of step 3 of the devised six steps, as described by[1]. For each bundle, we manually created an inclusion and exclusion waypoint ROI for both hemispheres. These waypoint ROIs were defined in MNI152 space and then transformed to the FOD template using FSL's FLIRT. Finally, for tracking, we used the *tckgen iFOD2* algorithm, which is capable of reconstructing fibers with complex configurations[5]. We used the following additional *tckgen* settings and inputs: step size 0.8mm, min. length = 8mm, max. length = 250 mm, max. number of streamlines = 10,000, unidirectional, include = inclusion ROI, exclude = exclusion ROI and seeding and cropped at GM-WM interface, with latter crop streamline more precisely as they cross WM-GM interface.

**S1.2. Fiber tracts refinement and cleaning**

The segmented tracts using our manually created waypoint ROIs were refined by comparing each candidate fiber bundle (i.e., ILF and Unc) to their respective probability maps provided within the AFQ tool[6]. As part of the AFQ processing, the probability maps are transformed into FOD template space. The ILF and Unc are assigned scores based on the probability values of the voxels through which they pass. Any trajectories with low probability scores are discarded. Finally, the selected ILF and Unc bundles should pass through the two predefined AFQ waypoint ROIs and also conform to the shape of the respective tract's probability map. In addition, the fiber tracts are cleaned further by determining the core of the fiber tracts to identify and remove any stray fibers. A fiber is represented as a 3D Gaussian distribution, and any outliers in the distribution are discarded (see Figure. S1 for an example of the processed tracts).

#### S1.3. Sampling diffusion measurements along WM tract lengths

The diffusion measurements were sampled along the ILF and Unc fiber cores at 100 equidistant points, which provided the respective tract profiles for each DKI map (MK, AK, RK, Kfa, MD, FA, and AWF) from the individual subjects, and from which the diffusion status in the WM can be inferred. Furthermore, to verify that diffusion profiling was restricted to WM and not contaminated with the signal originating from GM, we calculated the signed distance to the WM-GM boundary for each WM voxel, with negative values indicating proper sampling. For group-wise statistical analyses, each patient group was separated according to the side of lesion for MRI+ subjects, and the side of seizure focus for MRI- subjects (i.e., separated according to side ipsilateral to the epileptogenic temporal lobe), and their AFQ results were used as input to the Permutation Analysis of Linear Models (PALM) toolbox. In addition, geometrical properties of the tract profiles were represented using the first four statistical moments: mean (i.e., mean of quantitative values, mean(y) and center of gravity in the x-direction, mean(x)), standard deviation (SD), skewness (skew) and kurtosis (kurt)[7].

**Supplementary Figure**

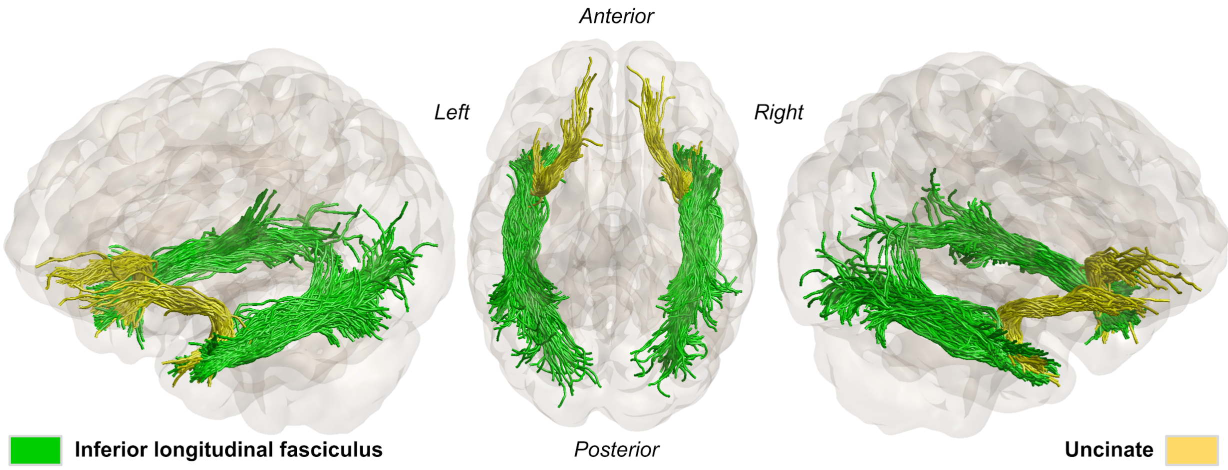


**Figure S1**. Showing the two WM bundles of interest, inferior longitudinal (green) and uncinate fasciculus (yellow) for a representative healthy subject. Generated using anatomically constrained tractography.

5. Tournier JD, Calamante F, Connelly A. Improved probabilistic streamlines tractography by 2nd order integration over fibre orientation distributions. In: Proceedings of the

international society for magnetic resonance in medicine [Internet]. Ismrm; 2010. Available from: https://cds.ismrm.org/protected/10MProceedings/PDFfiles/1670_4298.PDF

6. Yeatman JD, Dougherty RF, Myall NJ, Wandell BA, Feldman HM. Tract profiles of white matter properties: automating fiber-tract quantification. PLoS One. 2012 Nov 14;7(11):e49790.

7. Amunts K, Schleicher A, Bürgel U, Mohlberg H, Uylings HB, Zilles K. Broca's region revisited: cytoarchitecture and intersubject variability. J Comp Neurol. 1999 Sep 20;412(2):319–41.
